# The effects of increasing renal perfusion pressure on renal hemodynamics, microcirculation, and oxygen metabolism in a septic shock model

**DOI:** 10.1101/2025.09.27.678948

**Authors:** Luanluan Li, Ruiqiang Zheng, Jiangquan Yu

## Abstract

**Background:** The optimal target for mean arterial pressure (MAP) during clinical management of septic shock remains a topic of debate. This study investigates how elevating MAP, and thereby increasing renal perfusion pressure (RPP, defined as MAP-CVP), influences renal hemodynamics, microcirculation, and oxygen metabolism in a septic shock model.

**Methods:** Sixteen New Zealand rabbits of either sex and similar weight were randomly assigned to sham group (n=8) and septic shock group (n=8). Renal blood flow (RBF), microcirculatory blood flow and velocity in the kidney, as well as tissue oxygen pressure in the renal cortex and medulla, were assessed at four distinct time points: baseline (T0), model establishment (T1), restoration of MAP to baseline levels (T2), and an increase in MAP by more than 15% above baseline (T3).

**Results:** Within the septic shock group, although MAP and consequent RPP were increased, RBF markedly decreased from 17.63±2.50 to 8.50±1.93 ml/min (p < 0.05). No significant changes in microcirculation blood flow or flow velocity were observed between T2 and T3. Tissue oxygen pressure in the renal cortex and medulla declined significantly from 21.20±1.80 to 17.14±1.72 (p < 0.05) and from37.74±4.85 to 24.34±3.74 (p < 0.05), respectively.

**Conclusion:** Elevating RPP does not improve RBF, renal microcirculation, or tissue oxygen pressure in the renal cortex or medulla in septic shock. Elevating RPP does not improve RBF, renal microcirculation, or tissue oxygen pressure in the renal cortex or medulla in septic shock. We advocate restoring MAP to the baseline level rather than increasing it further to enhance renal perfusion pressure.

## Introduction

It is well known that sepsis is organ dysfunction caused by infection. Septic shock is a subset of sepsis [1].The 2021 international guidelines state that the treatment goal for patients with septic shock is a mean arterial pressure (MAP) ≥ 65 mmHg. A higher MAP does not improve the survival rate of patients with septic shock. Norepinephrine is the first-line drug for increasing blood pressure in septic shock. Therefore, we chose norepinephrine as the vasoconstrictor[2, 3] .By increasing the dose of norepinephrine to increase MAP, the renal perfusion pressure is increased. We use MAP-CVP to represent renal perfusion pressure[4, 5].An increase in MAP, a principal determinant of systemic circulation, directly results in an elevated RPP. Based on this foundation, whether elevating renal perfusion pressure can enhance renal blood flow, improve renal microcirculation, and optimize oxygen metabolism has become a critical focus of this experiment[6].A retrospective analysis of the MIMIC-IV database revealed that patients in the high mean perfusion pressure(MPP)group exhibited a higher in-hospital survival rate compared to those in the low MPP group [4].Nevertheless,conflicting evidence exists regarding the association between perfusion pressure, particularly as represented by MAP-CVP, and acute kidney injury (AKI) outcomes in critically ill patients. While some studies have found no significant association between platform pressure and either AKI incidence or recovery, others suggest that both excessively high and low perfusion pressures are linked to higher AKI incidence and lower recovery rates[7].Consequently, the optimal level of perfusion pressure in AKI patients remains a subject of ongoing debate. Moreover, there are limited animal studies directly elucidating the impact of increased MAP—aimed at augmenting RPP—on renal blood flow, microcirculation, and tissue oxygen metabolism in septic shock[8].Thus, the present experiment addresses an important gap in the research landscape[9].

## Method

### Surgical preparation

All rabbits underwent identical surgical protocols. Intramuscular injection of 25mg/kg ketamine [10].After anesthesia induction in the animals, ketamine was continuously pump into the vein at the ear margin[11].5% lidocaine was used for local anesthesia of the neck of New Zealand rabbits [12].A longitudinal incision was made 1cm below the thyroid cartilage to expose the trachea. A small incision was made laterally, and then an upward incision was made longitudinally (i.e., inverted T-incision of the trachea) [13-15].A 4F tracheal intubation was inserted for 3 to 4 cm. Connect to the ventilator for assisted ventilation and suture the incision. Ventilator parameters: VT: 24ml (6 ml/kg) PIP: 12-15cmH2O; PEEP: 3cmH2O;RR: 30 times/min; Flow: 6 L/min;FiO2: 35%, which can adjust the respiratory rate to maintain:PaO2:140-200mmHg and PaCO2:35-45mmHg[16].After successful tracheal intubation, midazolam, remifentanil and cisbenzene were pumped in to maintain anesthesia[17].A 4F central venous catheter was inserted into the right jugular vein, and a 3F Picco catheter was inserted into the left femoral artery and connected to the Picco machine. Blood pressure and CVP were measured, and MAP was calculated. The bilateral kidneys were accessed using the aforementioned method described in the literature [18].The renal arteries were located and separated. The renal blood flow probe was placed around the renal arteries, and the microcirculation probe was attached to the surface of the kidneys to detect the renal blood flow, microcirculation blood flow and velocity[14, 19].A sinus tract was drilled respectively in the renal cortex and medulla with a self-made needle. The optical fiber probe was inserted into the renal cortex and medulla to measure the tissue oxygen pressure in the renal cortex and medulla[3].After the rabbits were sacrificed after intravenous infusion of potassium chloride, the kidneys were removed to determine whether the sinus tract was in the renal cortex and medulla, see Fig.1[3, 20, 21].

**Fig. 1.**
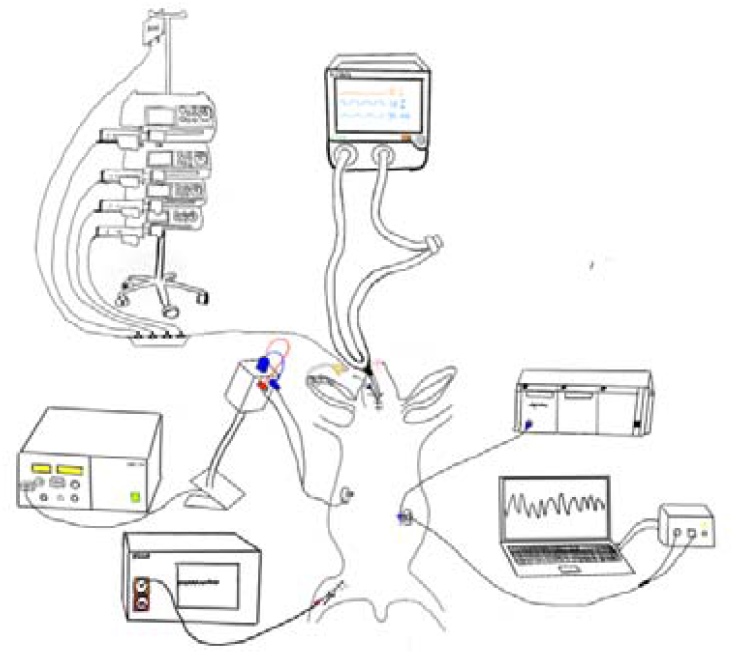
surgical preparation.

#### Experimental Protocol

Sixteen New Zealand white rabbits with similar body weights were randomly assigned to the sham group and the septic shock group.All animals received the aforementioned surgical preparation and were administered 50 ml/kg of 0.9% sodium chloride solution (normal saline) to meet physiological needs [3].The sham group received no further intervention.After the surgical preparation, when the condition of New Zealand white rabbits was stable, baseline measurements of various observation indicators were conducted and recorded as the T0 time point. Then, for the septic shock group,1.5 mg/kg LPS was injected via the ear vein [11, 15, 22-24].When the blood pressure of New Zealand white rabbits decreased to below 25% of the baseline value and was maintained for 30 minutes, it was considered that the model was successful[12, 23, 25].At this time, measurements at this point were recorded as T1. Immediately, a fluid infusion of 20 mg/kg and norepinephrine were administered to maintain MAP at baseline levels for one hour, at which time parameters were re-measured and denoted as T2. Subsequently, the dose of norepinephrine was increased to maintain MAP above 15% of the baseline value for one hour, with final measurements recorded as T3, see Fig.2.At the T0 time point in both groups, arterial blood was drawn from the ductus arteriosus and analyzed for blood gas parameters to confirm that the rabbits were in a normal physiological state. Following this, blood gas analysis was not repeated in the sham group, whereas it was performed at each time point in the septic shock group to assess the development of septic shock in the rabbits.

**Fig. 2.**
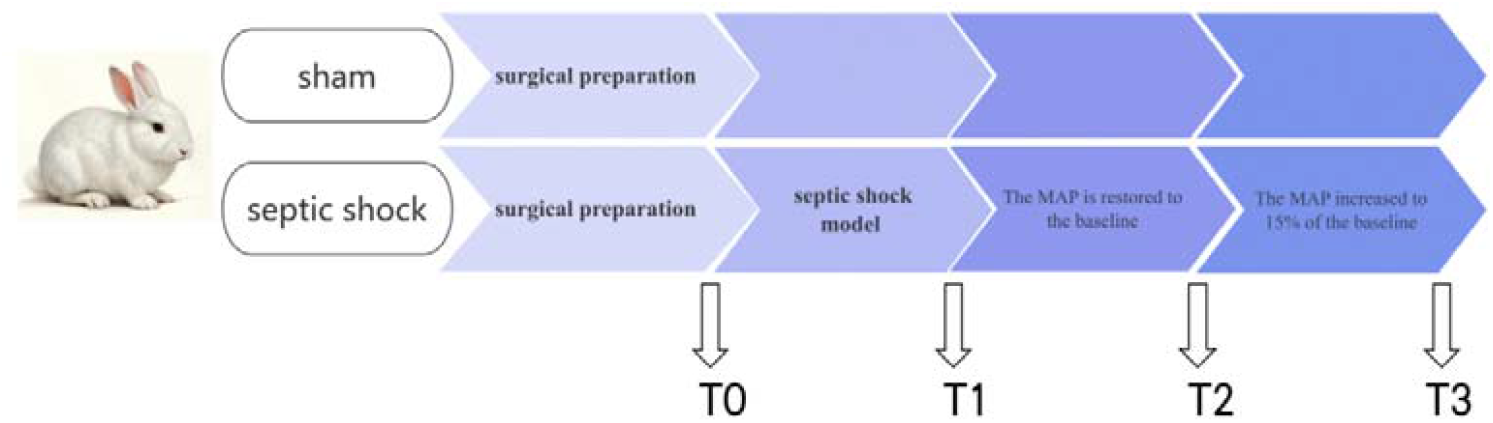
experimental protocol.

### Statistical analysis

The experimental data were analyzed using SPSS 29.0.1.0. After the experimental data were normally distributed, the quantitative data were expressed as mean ± standard deviation (X±SD) [14].Repeated measures analysis of variance was used to compare the differences between the two groups at four time points T0, T1, T2, and T3. Statistical significance was set at P<0.05. Additionally, differences among time points within each group were evaluated using Bonferroni correction to control for type I error[3].P<0.05 showed a statistically significant difference.Data visualization was conducted using Origin Pro 2024. Statistical significance is indicated as *p<0.05.

## Result

Table 1 shows the analysis of differences between and within groups at each time point between the two groups. There was no difference in MAP at each time point in the sham group. In the septic shock group, there was a difference between T0 and T1, no difference with T2, and a difference between T2 and T3, which was consistent with our experimental procedure, see Fig A. No significant differences in heart rate were detected, either within or between groups, across time points, though the septic shock group demonstrated a downward trend in heart rate,see Fig B. CVP remained stable in the sham group, whereas in the septic shock group, CVP decreased after model induction, although this change was not statistically significant compared to baseline. Restoration of MAP through fluid resuscitation resulted in a significant increase in CVP; however, inter-group differences remained non-significant, likely due to standardized physiological support provided to all animals,see Fig C. The RPP graph illustrates that renal perfusion pressure increased commensurately with MAP elevation,and this analysis is consistent with the MAP findings, matching the experimental design,see Fig D. Regarding RBF, significant differences were found between groups at T1, T2, and T3. No significant intra-group changes were found in RBF among sham group, but significant differences were seen in the septic shock group at all time points. Renal blood flow decreased after modeling and did not recover after restoring MAP, see Fig E. Microcirculation blood flow showed no significant difference at each time point in the sham group. In the septic shock group, it decreased significantly after modeling. No significant improvement in microcirculation was observed after the restoration of MAP, and no significant improvement was still observed after the increase of MAP. There was no statistically significant difference in microcirculation blood flow between groups T1, T2 and T3.As we can see from Table 1, there is almost no difference in microcirculation flow velocity between T1 and T2. The flow velocity at T3 begins to decrease, but there is no statistical difference, see Fig F and Fig G. The trends of tissue oxygen pressure in renal cortex and medulla were consistent. It decreased at each time point within both groups, and there were statistically significant differences. Our explanation was that measuring the oxygen pressure in tissues was an invasive operation, so the oxygen in tissues would also decrease to varying degrees during sham surgery. There were significant differences between the two groups at time points T1, T2, and T3. The pressure of oxygen in the tissue did not improve or stopped further deterioration, indicating that increasing MAP and increasing renal perfusion pressure could not improve oxygen metabolism, Fig H and Fig I.

**Table 1:** Analysis of differences in various observation indicators.

|  | 组别 | T0 | T1 | T2 | T3 |
| --- | --- | --- | --- | --- | --- |
| MAP (mmHg) | sham | 84.71 ± 2.51 | 87.58 ± 6.68# | 86.33 ± 2.22 | 84.17 ± 3.86+ |
|  | septic shock | 84.37 ± 3.86 | 58.54 ± 5.00a | 85.75 ± 3.48d | 98.79 ± 4.89cef |
| HR (bpm) | sham | 216.25 ± 11.97 | 208.00 ± 9.86 | 211.75 ± 11.34 | 205.38 ± 7.09 |
|  | septic shock | 217.38 ± 12.85 | 216.25 ± 12.67 | 208.00 ± 13.64 | 202.75 ± 15.63 |
| CVP (mmHg) | sham | 5.25 ± 0.71 | 5.38 ± 1.06 | 6.00 ± 1.51 | 5.88 ± 0.99 |
|  | septic shock | 5.38 ± 0.74 | 4.75 ± 1.04 | 7.13 ± 1.25bd | 6.13 ± 0.64 |
| RPP (mmHg) | sham | 79.46 ± 2.90 | 82.21 ± 5.95# | 80.33 ± 2.23 | 78.29 ± 4.05+ |
|  | septic shock | 79.00 ± 3.92 | 53.79 ± 4.96a | 78.62 ± 4.02d | 92.67 ± 4.66cef |
| RBF (ml/min) | sham | 43.63 ± 3.07 | 42.00 ± 2.56# | 41.38 ± 3.66* | 40.75 ± 2.82+ |
|  | septic shock | 41.50 ± 3.21 | 22.50 ± 2.78a | 17.63 ± 2.50bd | 8.50 ± 1.93cef |
| Microcirculation Flow (ml/min/100g) | sham | 41.22 ± 2.42 | 40.58 ± 2.34# | 41.14 ± 1.69* | 40.16 ± 2.33+ |
|  | septic shock | 40.68 ± 2.43 | 22.50 ± 1.68a | 21.70 ± 2.44b | 21.43 ± 1.73c |
| Microcirculation VEL (mm/sec) | sham | 1.92 ± 0.44 | 1.98 ± 0.41# | 2.10 ± 0.45* | 1.71 ± 0.45+ |
|  | septic shock | 1.93 ± 0.30 | 1.30 ± 0.10a | 1.31 ± 0.11b | 1.17 ± 0.08c |
| Cortical Tissue PO2 (mmHg) | sham | 37.31 ± 4.04 | 34.09 ± 3.64#a | 31.09 ± 4.00*bd | 27.30 ± 3.01+cef |
|  | septic shock | 38.28 ± 3.83 | 24.59 ± 3.60a | 21.20 ± 1.80bd | 17.14 ± 1.72cef |
| Medulla Tissue PO2 (mmHg) | sham | 60.00 ± 5.62 | 55.83 ± 6.12 | 53.13 ± 6.24 | 49.23 ± 5.78 |
|  | septic shock | 61.06 ± 4.85 | 47.59 ± 7.20 | 37.74 ± 4.85 | 24.34 ± 3.74 |

Notes: Differences at every time points within two group are denoted by "a, b, c, d", a-T0 vs T1, p<0.05; b-T0 vs T2, p<0.05; c-T0 vs T3, p<0.05; d-T1 vs T2, p<0.05; e-T1 vs T3, p<0.05; f-T2 vs T3, p<0.05. Between groups, differences at the four time points T0, T1, T2, and T3 are denoted by "\$, #, \*, +" respectively.

## Discussion

In the septic shock model, increases in mean arterial pressure (MAP)—a surrogate for systemic circulation—result in higher renal perfusion pressure (RPP) [26].We can observe that the renal blood flow does not increase, indicating that the increase in MAP does not lead to an increase in renal blood flow, nor does it improve microcirculation, nor does it increase oxygen metabolism. That is to say, the increase in perfusion pressure has no linear relationship with the increase in renal blood flow, microcirculation and the improvement of oxygen metabolism. Notably, at T3, with MAP increased to more than 15% above baseline, RBF decreased while renal microcirculation remained unchanged. This raises the question of whether the reduction in renal blood flow during elevated MAP in septic shock may inorder to preserve microcirculatory perfusion[27, 28].Possible mechanisms may include intrinsic renal autoregulation, protective responses to elevated MAP, or pharmacologic effects of norepinephrine[29, 30].These hypotheses warrant additional investigation for clarification.At the same time, it should be emphasized that raising MAP to normal levels is not without clinical significance in rabbits with septic shock. While elevating blood pressure to the normal range undoubtedly provides physiological benefits during the management of septic shock, this intervention alone does not lead to a reduction in lactic acid levels. The core pathophysiological issue in septic shock remains uncontrolled infection; therefore, etiological treatment is essential in clinical practice. Merely correcting hypotension is insufficient for effective management.

### Limitation

In the septic shock group, the lactate level of New Zealand rabbits was basically greater than 2mmol/L at the modeling time point. Meanwhile, when MAP increased by 15% above baseline and was sustained for one hour, lactate levels continued to rise, suggesting ongoing progression of the septic shock model, see supplementary materials.Beyond that, at the T3 time point, ongoing decreases in RBF imply that the septic shock model might still be actively evolving, and it is not possible to definitively conclude that elevated perfusion pressure is harmful to renal blood flow[29].

Rather, it indicates that elevated renal perfusion pressure does not necessarily enhance renal blood flow. In fact, there is currently insufficient evidence to confirm that an increase in “renal perfusion pressure”-something is really which waiting for us to find does’t result in improvements in renal blood flow, microcirculation, or oxygen metabolism. However, it is certain that when RPP is defined solely as MAP-CVP, these parameters—renal blood flow, microcirculation, and oxygen metabolism —do not consistently improve[7, 31].In other words, can MAP-CVP truly reflect “renal perfusion pressure”?

## Conclusion

In the septic shock model, by increasing MAP to raise renal perfusion pressure, we found it cannot increase renal blood flow, improve microcirculation, or increase tissue oxygen pressure in renal cortex and medulla[32].We more advocate increasing the MAP to the baseline. There is no need for a higher MAP to increase the renal perfusion pressure[29, 33].

## Abbreviations

MAP: Mean Arterial Pressure
RPP: Renal Perfusion Pressure
RBF: Renal Blood Flow
CVP: Central Venous Pressure

## Acknowledgements

Specially, thanks to Yunfan Feng, Tianwei Wang and Yuanjin Pan for their strong support in the pre-experiment. Lu Xu is in charge of material procurement.Yuchen Wang raised the rabbits in the pre-experiment for a week.

## Author contributions

Luanluan Li is responsible for project design, data collection, data analysis, article writing, diagram drawing and animal breeding ; Ruiqiang Zheng is in charge of the project funds and experimental guidance;Ruiqiang Zheng and Jiangquan Yu is responsible for the advance payment of equipment funds.

## Availability of data and materials

The datasets used and/or analyzed during the current study are available from the corresponding author upon reasonable request.

## Declarations

### Ethics approval and consent to participate

The experimental procedures were reviewed and approved by the Animal Ethics Committee of Medical College of Yangzhou University.

### Consent for publication

Not applicable.

---

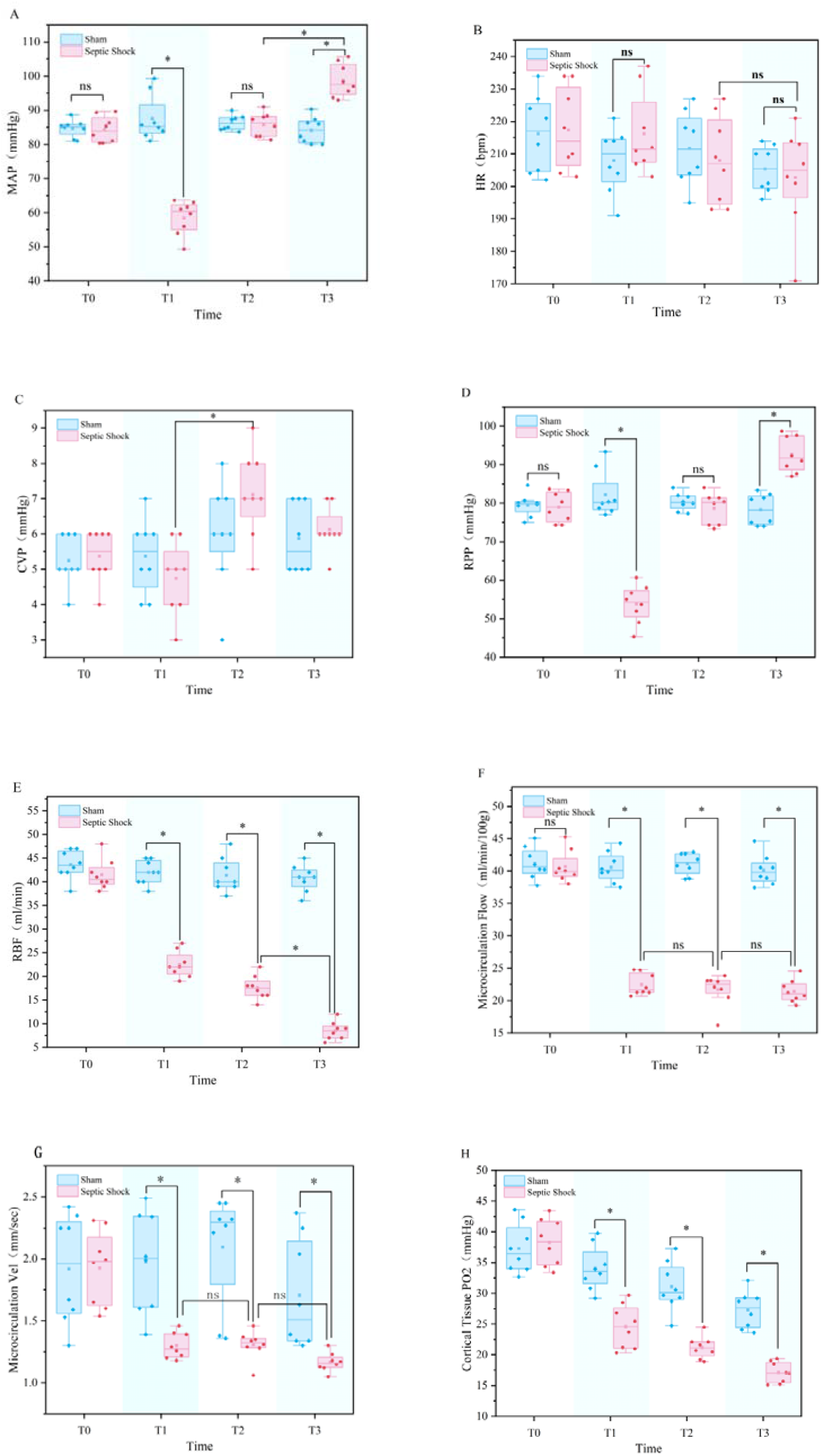
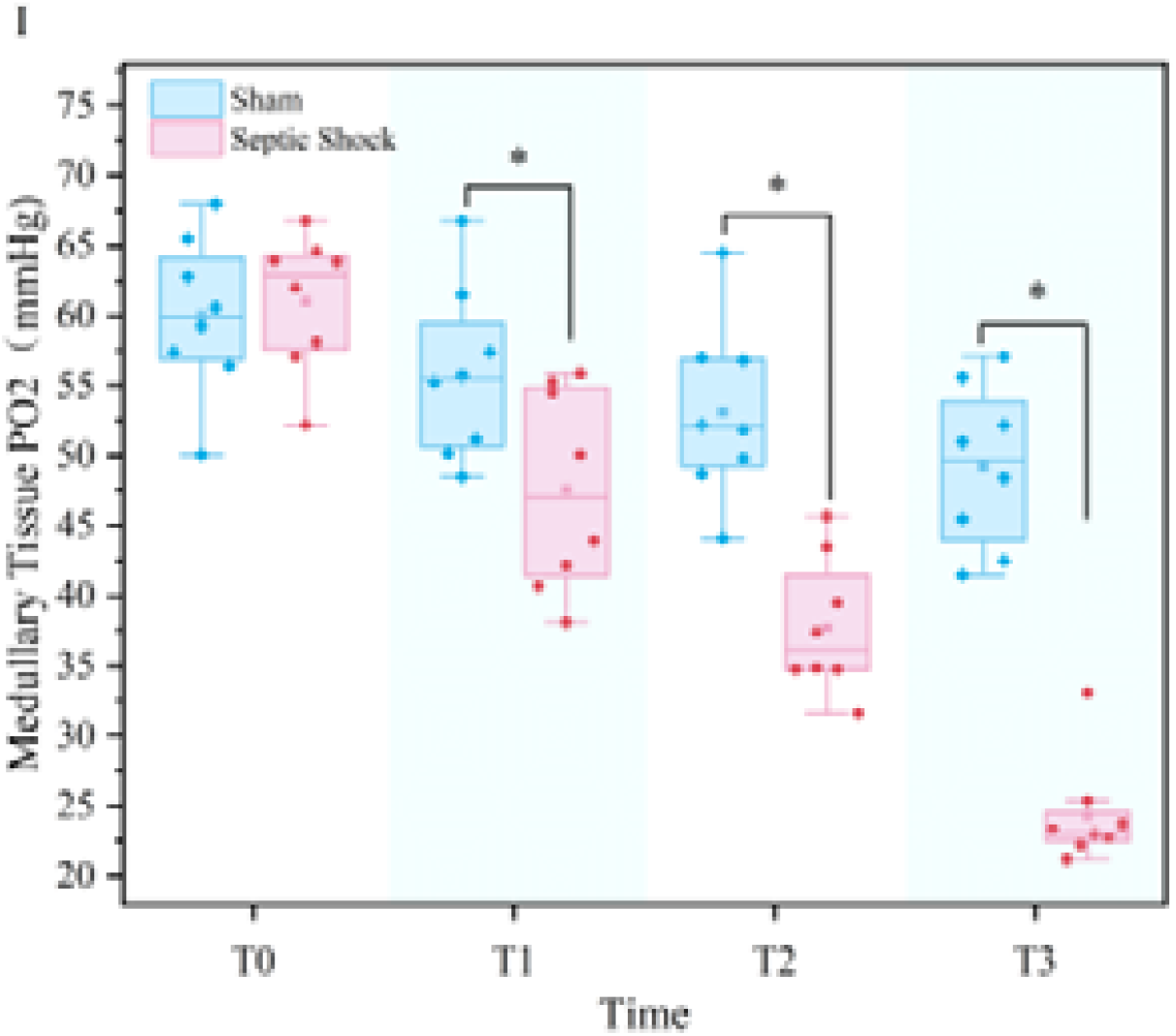

